# Macrophage-Based iNos Reporter Reveals Polarization and Reprogramming in the Context of Breast Cancer

**DOI:** 10.1101/2022.08.18.504212

**Authors:** Javier A. Mas-Rosario, Josue D. Medor, Mary I. Jeffway, José M. Martínez-Montes, Michelle E. Farkas

## Abstract

As part of the first line of defense against pathogens, macrophages possess the ability to differentiate into divergent phenotypes with varying functions. The process by which these cells change their characteristics, commonly referred to as macrophage polarization, allows them to change into broadly pro-inflammatory (M1) or anti-inflammatory (M2) subtypes, and depends on the polarizing stimuli. Deregulation of macrophage phenotypes can result in different pathologies or affect the nature of some diseases, such as cancer and atherosclerosis. For this reason, it is necessary to better understand macrophage phenotype conversion in relevant models. However, there are few existing probes to track macrophage changes in multicellular environments. In this study, we generated an eGFP reporter cell line based on inducible nitric oxide synthase (*iNos*) promoter activity in RAW264.7 cells (RAW:*iNos*-eGFP). iNos is associated with macrophage activation to pro-inflammatory states, and decreases in immune-suppressing ones. We validated the fidelity of the reporter for *iNos*, including following cytokine-mediated polarization, and confirmed that reporter and parental cells behaved similarly. RAW:*iNos*-eGFP cells were then used to track macrophage responses in different *in vitro* breast cancer models, and their re-education from anti- to pro-inflammatory phenotypes via a previously reported pyrimido(5,4-b)indole small molecule, PBI1. Using two mouse mammary carcinoma cell lines, 4T1 and EMT6, effects on macrophages were assessed via conditioned media, two-dimensional/monolayer co-culture, and three-dimensional spheroid models. While conditioned media derived from 4T1 or EMT6 cells and monolayer co-cultures of each with RAW:*iNos*-eGFP cells all resulted in decreased fluorescence, the trends and extents of effects differed. We also observed decreases in *iNos*-eGFP signal in the macrophages in co-culture assays with 4T1- or EMT6-based spheroids. We then showed that we are able to enhance *iNos* production in the context of these cancer models using PBI1, tracking increased fluorescence. Taken together, we demonstrate that this reporter-based approach provides a facile means to study macrophage responses in complex, multicomponent environments. Beyond the initial studies presented here, this platform can be used with a variety of *in vitro* models and extended to *in vivo* applications with intravital imaging.

## 1. Introduction

Macrophages are cells of the innate immune system that play important roles in fighting infections and supporting tissue development, maintenance, and remodeling.^1,2^ They reside in tissues, body cavities, and mucosal surfaces (including but not limited to the lungs, spleen, skin, heart, kidney, and peritoneum), and contribute to both homeostasis and disease.^3^ These cells are “plastic,” which refers to their capacity to alter their phenotypes in a process known as macrophage polarization (**Figure S1**).^4,5^ This process is dictated by surrounding pathogens or cytokines that influence macrophage phenotypes and responses.

Macrophages can respond to both innate and foreign pro-inflammatory signals, including cytokines, such as interferon gamma (IFN-γ) and tumor necrosis factor alpha (TNF-α), or lipopolysaccharides (LPS), respectively. These result in what are referred as immune-stimulating, or classically activated macrophage phenotypes, commonly referred to as M1.^4^ This subtype of macrophages has pathogen-killing abilities and has been shown to eliminate tumor cells via enhanced phagocytosis and generation of reactive oxygen and nitrogen species (ROS and RNS, respectively).^6^ Previous studies in murine primary macrophages showed that M1 macrophages are characterized by enhanced expression of toll-like receptor 2 (Tlr-2),^7^ intracellular adhesion molecule 1 (Icam1),^8^ Tnf-α,^9^ and inducible nitric oxide (iNos),^10^ and have decreased expression of mannose receptor (MR/Cd206),^11^ early growth response protein 2 (Egr2/Krox20),^12,13^ cluster of differentiation 36 (Cd36)^14^ and neuropilin 1 (Nrp1).^15^

At the other end of the spectrum, macrophages can assume roles associated with immune suppression and wound-healing responses. When macrophages are stimulated with interleukin 4 (IL-4), interleukin 10 (IL-10), interleukin 13 (IL-13), or other anti-inflammatory cytokines (described further below), they adopt an immune-suppressive M2 subtype.^16^ Compared to the M1 phenotype, these have been shown to exhibit opposite patterns of expression of the aforementioned polarization markers. Due to their complexity, M2 macrophages may be further classified into M2a, M2b, M2c, and M2d/tumor-associated macrophage (TAM) categories. Each results from the presence of specific cytokines, and while some common characteristics are shared, others are unique.^5,17^

Macrophages can be re-educated from one phenotype to another when the conditions in their surrounding environments change.^18^ In the case of cancer, tumor cells possess the ability to secrete anti-inflammatory cytokines, including IL-4, IL-10, IL-13, and others,^19^ that can convert undifferentiated (M0) or M1 macrophages into the M2d/TAM phenotype.^10,12^ TAMs have been shown to aid in multiple aspects of cancer, including tumor growth, angiogenesis, remodeling of the tumor microenvironment, invasion, and establishment and maintenance of metastases.^19^ Macrophages have been shown to be involved in multiple types of cancer, including breast, lung, gastric, colorectal, and pancreatic cancers, with these cells sometimes contributing up to 50% of the tumor mass.^20^ Studies have also shown that high infiltration of TAMs in tumor tissues are correlated with poor patient prognoses.^21,22^ Due to their implications in cancer and potential to act against it, macrophage re-programming is of interest in immunotherapeutic treatments.^6,23^ Therefore, it is important to gain a better understanding of the interactions between cancers and macrophages, and their outcomes.

To further elucidate their role in the context of cancer, various techniques have been used for tracking macrophage behavior in *in vitro* and *in vivo* cancer models. Real Time Polymerase Chain Reaction (RT-PCR), for instance, is a technique used to identify macrophage phenotypes *in vitro* based on the expression of phenotype-associated markers. However, this technique is labor intensive and expensive, it requires use or isolation of a single cell type, typically provides average mRNA expression levels, and is difficult to obtain multiple time-points with.^24^ Enzyme-linked immunosorbent assays (ELISAs), are similarly time-consuming and expensive, and require single cell types to be able to assess which cells produce given markers, resulting in population-level data.^25^ Immuno-staining^26^ and flow cytometry^27^ are both able to provide data for individual cells, but like RT-PCR, are limited to evaluations at single time points, and macrophage-specific markers must be used to differentiate them from other cell types. In terms of *in vivo* approaches, some of the most commonly used methods for tracking macrophages in animal models are bioluminescence imaging (BLI) and intravital microscopy, which require fluorescent probes for macrophage labeling (e.g., intrinsic fluorophores (NADH, FAD, and collagen), genetic probes (fluorescent proteins), and commercial chemical probes (fluorescent dye-conjugates)).^28^ These techniques facilitate the tracking of macrophages *in vivo*, but fail to show phenotypic changes in the cells. Therefore, there is a need for methods that allow the tracking of macrophages in multi-cellular environments and the visualization of changes in their phenotype in real time.

Various platforms have been used to study the effects of cancers on macrophages. While some are more physiologically relevant than others, the mode of assessing phenotypic markers is often the dictating factor. The most commonly employed models include exposing macrophages to cancer cell-derived conditioned media,^29^ co-culturing them with cancer cells in a monolayer,^30^ or more complex experimental designs, such as tumor spheroids or other *in vitro* 3D models.^31^ Conditioned media refers to a collection of secreted signaling proteins (secretome) from cells of interest, and is commonly used to study the effects of cancer on macrophages and other immune cells.^32-35^ While it is compatible with most of the techniques described above (since only a single population of cells is present), it excludes cell-to-cell interactions, which play key roles in diseases like cancer. Macrophages and other cell types are able to associate and influence one another in both two- and three-dimensional, or 2D and 3D, co-cultures. In the 2D model, cells grow in a monolayer, typically attached to a plastic surface.^36-38^ This method is useful for studying cell-to-cell interactions, simple to maintain, and amenable to functional tests (e.g., phagocytosis of cancer cells by co-cultured immune cells).^39^ However, 2D cultures have limitations that include alterations in cell morphology, polarity, and method of division.^37,38^ 3D cultures, where cells grow in three dimensions,^40^ better represent cell-to-cell and cell-to-extracellular environment interactions, morphology and cell division, and access to oxygen, nutrients, metabolites, and signaling molecules or cytokines.^37,41,42^ The use of 3D co-cultures is also highly relevant, since the characteristics of cells and their responses, including to drugs, can be differ based on whether they are cultured in two versus three dimensions.^42^ In both cases, however, while the cellular environments are more realistic, the means of assessing macrophage polarization becomes more difficult. For this reason, we generated a reporter cell line to track the expression of a phenotype-associated marker in relevant disease models over time and following drug treatment.

In this work, we generated a fluorescent macrophage phenotype reporter cell line (RAW:*iNos*-eGFP) based on polarization-associated marker iNos. We confirmed the fidelity of the RAW:*iNos*-eGFP cells, including following exposure to polarizing cytokines, and established that reporter and parental cells behaved similarly. The reporter cells were then used to monitor macrophage responses in different *in vitro* breast cancer models, and their re-education from anti- to pro-inflammatory phenotypes via a previously reported Tlr4-agonist, PBI1.^6^ Effects of 4T1 and EMT6 cell lines on macrophages were assessed via conditioned media, two-dimensional/monolayer co-culture, and three-dimensional spheroid models. While conditioned media derived from 4T1 or EMT6 cells and monolayer co-cultures of each with RAW:*iNos*-eGFP cells resulted in decreased fluorescence, the trends and extents of effects differed. We also observed a decrease in *iNos*-eGFP signal in the macrophages in 3D culture assays with 4T1- or EMT6-based spheroids. We then show that we are able to induce an increase in *iNos* production, even in the presence of 3D, M2-polarizing cancer models using PBI1. Taken together, we demonstrate that this reporter-based approach provides an easier and more efficient means to study macrophage responses in more relevant and complex, multicomponent environments. Our preliminary results, show that this platform has the potential of being used with a variety of *in vitro* models and extended to *in vivo* applications.

## 2. Materials and Methods

### Cell culture

Murine RAW264.7 macrophages and 4T1 and EMT6 murine mammary carcinoma cells were purchased from American Type Culture Collection (ATCC). Human embryonic kidney (HEK293) cells were obtained from Prof. D. Joseph Jerry (Veterinary and Animal Sciences, UMass Amherst). All cell lines, including RAW:*iNos*-eGFP, were cultured at 37 °C under a humidified atmosphere containing 5% CO_2_. Standard growth media consisted of high glucose Dulbecco’s Modified Eagle Medium (DMEM, Gibco) supplemented with 10% fetal bovine serum (FBS, Corning), 1% L-Glutamine (200 mM, Gibco) and 1% antibiotics (100 μg/ml penicillin and 100 μg/ml streptomycin, Gibco) – herein referred to as complete DMEM. Under the above culture conditions the cells were sub-cultured approximately once every 3-4 days and only cells between passages 7 and 20 were used for all experiments.

### Macrophage polarization

To polarize macrophages for sorting, cells were plated in a T75 culture flask and grown at 37 °C under a humidified atmosphere containing 5% CO_2_ to confluence prior to polarization. Once confluent, culture media was removed and replaced with media containing either 50 ng/mL interleukin 4 (IL-4) (BioLegend) for 48 hr to generate M2 macrophages or 50 ng/mL interferon-gamma (IFN-γ) (BD Biosciences) and 50 ng/mL of lipopolysaccharide (LPS) (Sigma Aldrich) for 24 h for M1 macrophages. For confocal microscopy and RT-PCR experiments, cells were plated in three biological replicates in an 8-well Lab-Tek II chambered cover-glass system plate (Nunc) or 24-well plates at a density of 100,000 cells/well in 500 µL of media and incubated at 37 °C under a humidified atmosphere containing 5% CO_2_. Non-treated macrophages (M0; grown in complete DMEM only) were used as controls. To generate M2 macrophages, 24 h after plating, the media was removed and replaced with complete DMEM media containing 50 ng/mL IL-4 and incubated for an additional 48 h at 37 °C under a humidified atmosphere containing 5% CO_2_. To generate M1 macrophages, culture media was removed and replaced with complete DMEM containing 50 ng/mL IFN-γ and 50 ng/mL of LPS 48 h after plating and incubated for an additional 24 h. 72 h after plating (48 h after treatment with M2 cytokines and 24 h after treatment with M1 cytokines), cells were used further in experiments as indicated.

### RNA extraction and cDNA conversion

Cells were lysed, and approximately 1.5 μg RNA was harvested from each well using the PureLink RNA Mini Kit (Ambion) following the manufacturer’s instructions. To convert RNA to complementary DNA (cDNA), 1 µL of 50 μM random hexamers (Applied Biosystems) and 1 µL of 10 mM dNTPs (Thermo Scientific) were added to 11 µL of RNA and heated at 65 °C for 5 min for annealing. Then, 1 µL/sample of 40 U/μL RNaseOut (Invitrogen), 1 µL/sample of 200 U/μL SuperScript IV Reverse Transcriptase (Invitrogen), 1 µL/sample of 100 mM DTT (Invitrogen), and 4 µL/sample of 5x Super Script IV buffer (Invitrogen) were added and amplification proceeded at 53 °C for 10 min and melting at 80 °C for 10 min. The resulting cDNA was frozen at -20 °C and used for RT-PCR experiments within 1 week. RNA and cDNA were quantified using a NanoDrop 2000 (Thermo Fisher). RNA and cDNA contamination and integrity were assessed by analyzing the A260/A280 ratio, where ratios greater 1.8 for DNA and 2.0 for RNA were considered pure.

### Quantitative RT-PCR

RT-PCR was performed on the cDNA generated using a CFX Connect real-time system (Biorad) with iTaq Universal SYBR Green Supermix (Biorad). All DNA primers were purchased from Integrated DNA Technologies. The following primer sequences were used: β-actin (forward) 5’-GATCAGCAAGCAGGAGTACGA-3’, (reverse) 5’-AAAACGC-AGCGCAGTAACAGT-3’; iNos (forward) 5’-GTTCTCAGCCCAACAATACAAGA-3’, (reverse) 5’-GTGGACGGGTCGATGTCAC-3’. The reaction mixtures included 200 nM of each primer, 100 ng of cDNA, 10 μL SYBR green supermix, and H_2_O to a final volume of 20 μL. Analyses were performed as follows: the samples were first activated at 50 °C for 2 min, then 95 °C for 2 min. Denaturing occurred at 95 °C for 30 s followed by annealing at 58 °C; the denature/anneal process was repeated over 40 cycles. Relative gene expression was determined by comparing the Ct value of the gene of interest to that of the β-actin housekeeping gene, by the 2ΔΔCt method.^43^ Three biological replicates were performed for each treatment condition and three technical replicates were used for each biological replicate. Data was analyzed using CFX Manager 3.1 software (Biorad). Cq values were generated by using the point at which the sample fluorescence value exceeded the software’s default threshold value. Each sample was normalized to the non-treated control.

### Molecular cloning of *iNos*-eGFP lentiviral plasmid

To construct the lentiviral *iNos*-eGFP reporter construct, a plasmid containing the promoter region of Mus musculus iNos was obtained from Addgene (pGL2-NOS2 Promoter-Luciferase – Plasmid # 19296 from Charles Lowenstein).^44^ The following primers were designed and used to amplify the promoter for the gene of interest and incorporate XhoI and BamHI sites at the 5’ and 3’ ends, respectively, underlined): iNos-XhoI (forward) 5’-<u>CCGCTCGAGCGG</u>CGAGCTCTTACGCGGACTTT-3’ and iNos-BamHI (reverse) 5’-<u>CGCGGATCCGCG</u>TTTACCAACAGTACCGGAAT-3’. PCR was performed using Phusion High Fidelity Master Mix (NEB) using optimized conditions (higher temperatures for annealing/extension of 72 °C) due to the high GC content of the primers. Following purification, the resulting ∼1.3 kb fragment was subcloned into the pRRLSIN.cPPT.PGK-GFP.WPRE lentiviral vector (Addgene plasmid # 12252 from Didier Trono). Both the PCR product and the recipient plasmid were digested with XhoI and BamHI (NEB) according to manufacturer’s protocols, followed by purification. Ligations were performed using T4 ligase (NEB) according to manufacturer’s protocols. Ligation mixtures were then transformed into STBL3 bacteria (ThermoFisher) by electroporation and plated for overnight (approximately 18 h) growth in ampicillin-containing agar plates at 37 °C. Single colonies were then picked and transferred into 5 mL of LB media with ampicillin for further expansion in a shaker incubator at 37 °C. 12 h later, the 5 mL culture was diluted to 50 mL using LB media containing ampicillin, and returned to the incubator for overnight growth. Sanger sequencing was performed by GeneWiz to confirm the final construct.

### Generation of stable RAW:*iNos*-eGFP cells – Lentiviral transductions

HEK293T cells were seeded in 60 mm culture dishes and transiently transfected with 3 μg psPAX2 packaging plasmid, 2 μg pMD2G envelope plasmid (both from Prof. D. Joseph Jerry, Veterinary and Animal Sciences, UMass Amherst), and 5 µg *iNos*-eGFP reporter constructs generated above, using Lipofectamine3000 (ThermoFisher Scientific), according to the manufacturer’s instructions. Lentiviral particles were harvested from the supernatant 48 h after DNA-lipid complexes were added to cells. The virus-containing supernatant was passed through a 0.45 µm filter. Equal volumes of lentivirus-containing supernatant and complete DMEM containing 4 µg/mL polybrene (Sigma) were combined. Confluent RAW264.7 cells grown in T25 flasks were treated with 6 mL of lentivirus-containing media. Infections were performed every 12 h over 48 h (total of 4 infections), after which the medium was replaced with complete DMEM, and the cells were allowed to recover, grow, and expand for 2-3 days to ensure a viable population. Cells were then prepared for sorting of positive cells as described below.

### Cell Sorting of RAW:*iNos*-eGFP

To ensure a homogenous population, cells were sorted twice under different polarizing conditions. For the first sorting, cells were exposed to M1-polarizing cytokines for 24 h (described above) to induce an M1 phenotype. After 24 h, cells were detached from the cell culture flask; 5-7 × 10^6^ cells were resuspended in 3 mL of FACS buffer (4% FBS in phosphate buffered saline (PBS, Gibco)) and sorted at the University of Massachusetts Amherst Flow Cytometry Core Facility using a BD FACSAria Fusion (Becton Dickinson). The instrument was configured with 4 lasers (405 nm, 488 nm, 561 nm, 640 nm), and a 100 µm nozzle size was used for sorting. Of 2.18 × 10^6^ positive cells, the top 20.6% of the highest fluorescing cells (436,000 cells with ∼98% purity) were sorted. These were plated in T25 flasks for recovery, expansion, and further sorting. For the second sorting, cells were then treated with M2 cytokines for 48 h. Once polarized, 5-7 × 10^6^ cells were resuspended in FACS buffer and sorted using the same instrument and configuration from the first. Of 3.7 × 10^6^ positive cells, the bottom 10.5% of the lowest fluorescing cells (370,000 cells with ∼98% purity) were sorted and plated in T25 flasks for further expansion and use.

### Confocal microscopy

For acquisition of cell morphology and fluorescence images, RAW:*iNos*-eGFP cells were plated in a 8-well Lab-Tek II chambered cover-glass system plate (Nunc) at a density of 100,000 cells/mL in 500 µL (50,000 cells/well) and allowed to adhere overnight. All treatments of RAW:*iNos*-eGFP cells were performed 24 h after being plated. Cells were imaged every 24 h for up to 72 h depending on the experiment. After polarization as described above, cells were imaged using a Nikon Ti-E C2 confocal microscope at 10x magnification. ImageJ/Fiji software was used for the quantification of fluorescence of the confocal images on a per-cell basis via thresholding method.^45,46^ Every experimental group was reproduced with three biological replicates, for which a single image was acquired for each that included between ∼100-1000 cells for which the integrated fluorescence intensity (mean fluorescence X area of the cell) was defined.

### Generation of conditioned media

Cells were cultured and passaged at least once before being used to generate conditioned media. The procedure used to generate 4T1- and EMT6-conditioned media follows a previously established protocol.^47^ Briefly, cells were cultured in T175 flasks with complete DMEM until they became >90% confluent. At that point, the media was replaced with complete DMEM media and cells were cultured for an additional 7 days. On day 7, the media was collected and filtered through a 0.45 µm syringe filter and stored at -20 °C. For the experiments described here, it was used within the first six months.

### Treatment with 4T1 or EMT6 conditioned media

RAW:*iNos*-eGFP cells were plated in an 8-well Lab-Tek II chambered cover-glass system plate at a cell density of 50,000 cells/well in 500 µL complete DMEM and allowed to adhere overnight at 37 °C, 5% CO_2_. After 24 h, the culture media was removed and replaced with 40% conditioned media from 4T1 or EMT6 cells and 60% complete DMEM. Cells were incubated with conditioned media for 48 h at 37 °C, 5% CO_2_ and were then assessed via confocal microscopy.

### Spheroid generation

Using the hanging drop technique,^48^ 10 µL-droplets of a 10^6^ cell/mL-solution of 4T1 or EMT6 cells were added to a 60 mm petri dish lid, and 3-4 mL of PBS was added to the bottom of the petri dish. The lids were immediately inverted and placed atop the dishes containing the PBS reservoir. The cells were then incubated at 37 °C at 5% CO_2_ for 3-4 days to allow seeds to form. The seeds (typically between 5-15 per plate) were individually transferred from the 10 µL hanging drops using a 100 µL-pipette tip (with the tip cut off) to a 25 mL-round bottom flask containing 5 mL of complete DMEM. Flasks were placed on a platform shaker within a cell culture incubator and grown at 37 °C at 5% CO_2_ with shaking at 150 rpm for an additional 3-4 days. Spheroids were subsequently drawn from the flask and used directly in the respective experiments.

### Two- and three-dimensional co-cultures

For two-dimensional (2D) co-cultures, RAW:*iNos*-eGFP cells were concurrently plated with 4T1, and separately, EMT6 breast cancer cells at a 1:1 ratio, 250 µL of each for a total of 500 µL, in complete DMEM using an 8-well Lab-Tek II chambered cover-glass system plate. For three-dimensional (3D) co-cultures, 50,000 RAW:*iNos*-eGFP cells in 500 µL complete DMEM were plated in an 8-well Lab-Tek II chambered cover-glass system, and allowed to adhere overnight. 24 h after plating, a single spheroid in 20 µL complete DMEM was added to the macrophage monolayer using a 100 µL-pipet and imaged immediately following. Images were acquired every 24 h for up to 72 h for both 2D- and 3D co-culture experiments. Experiments were performed in biological triplicates.

### Small molecule (PBI1) treatment

For experiments involving treatment with pyrimido(5-4b)indole (PBI1, synthesized as previously described),^6^ a 5 mg/mL stock solution of PBI1 dissolved in dimethyl sulfoxide (DMSO; Sigma-Aldrich) was prepared. Cell treatments were then prepared by adding 2 µL of PBI1 stock solution per 500 µL of cell culture media for a final dosing concentration of 20 µg/mL of PBI1 and 0.4% DMSO. Media in 8-well Lab-Tek II chambered cover-glass plate containing cells was removed and replaced with 500 µL of PBI1-containing media (20 µg/mL) for 24 h prior to analysis.

## 3. Results and Discussion

### Generation and Validation of RAW:*iNos*-eGFP Cell Line

Because we sought to generate cell-based reporters that could be easily and consistently used, we elected to use immortalized cells for our system. RAW264.7 cells are an immortalized cell line, commonly used as a model for macrophages.^49^ We first confirmed the similarity of the candidate marker, *iNos*’s expression profiles between RAW264.7 and primary bone derived macrophages (BMDMs). Both cell types were assessed under non-polarized (M0), and immune-activating (M1) and –suppressing (M2) states. We found that while the ratios of *iNos* levels between phenotypes varied between the primary and immortalized cells, the trends of *iNos* being substantially higher for M1 and slightly lower for M2, respective to M0, were similar (**Figure S2**).

After verifying our marker for use as a reporter, molecular cloning methods were used to generate a lentiviral plasmid for enhanced green fluorescent protein (eGFP) expression under the control of the *iNos* promoter sequence. This construct was stably transfected into RAW264.7 cells to yield the RAW:*iNos*-eGFP reporter cell line. We anticipated that the reporter cell line would increase in fluorescence when adopting a pro-inflammatory M1 phenotype and become dimmer when assuming an anti-inflammatory M2 state. Following transfection and preliminary confirmation of fluorescence via microscopy (data not shown), cells were sequentially sorted via flow cytometry under polarizing conditions. First, transfected cells were polarized to the M1 phenotype via cytokine (LPS and IFN-γ) treatment, and the top 20% of the cells with the highest fluorescence levels were selected (**Figure S3, left**). Then, this population of cells was polarized to the M2 phenotype using IL-4, and the bottom 10% of the cells with the lowest fluorescence were selected (**Figure S3, right**).

The resulting RAW:*iNos*-eGFP cell line was evaluated via RT-PCR and confocal microscopy. In both cases, the reporter cells were polarized to M1 and M2 phenotypes (non-treated cells represented M0), and relative *iNos* and eGFP fluorescence levels were determined and compared. RT-PCR results indicated that the *iNos* expression patterns of RAW:*iNos*-eGFP (**Figure 1a**) were similar to those of the RAW264.7 parental cell line (**Figure S2**). This suggests that both *iNos* expression and macrophage polarization pathways were unaffected by the insertion of the plasmid construct. To confirm fluorescence changes resulting from the promoter-driven reporter, polarized and non-polarized cells were evaluated using confocal microscopy (**Figure 1b**). Significantly increased fluorescence was observed for cells in the M1 phenotype and diminished levels were observed for the M2 subtype, as determined by quantification on a per cell basis (**Figure 1c**).

**Figure 1:**
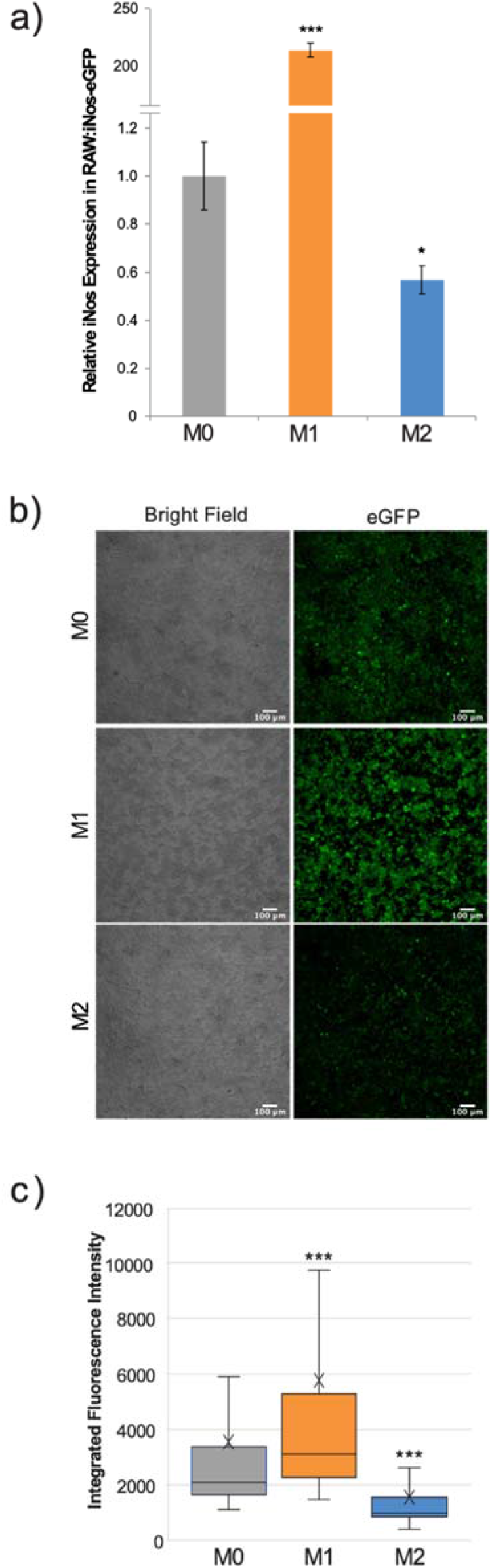
Effects of cytokine polarization on RAW:iNos-eGFP cells. (a) RT-PCR data showing relative levels of *iNos* mRNA across polarization states (M0 = gray, M1 = orange, and M2 = blue). Error bars represent standard error. (b) Confocal microscopy images acquired following polarization of RAW:*iNos*-eGFP. BF = bright field, eGFP = enhanced green fluorescent protein. Scale bars on confocal images represent 100 µm. (c) Quantification of fluorescence intensity on a per-cell basis from the figures shown on the left. M0 = gray (n = 1,710), M1 = orange (n = 1,820), and M2 = blue (n = 1,142). The “x” in the box represents the mean; the bottom and top lines of the box represent the median of the bottom half (1st quartile) and median of the top half (3rd quartile), respectively; the line in the middle of the box represents the median; the whiskers extend from the ends of the box to the minimum value and maximum value. For all panels, M0 = non-treated; M1 = LPS/IFN-γ; M2 = IL-4. For (a) and (c), Student T-test was used for statistical analysis comparing M1 and M2 macrophages to M0 (p<0.05 = *, p<0.001 = ***).

### RAW:*iNos*-eGFP Responses to Different Breast Cancer Models

Following validation, the RAW:*iNos*-eGFP reporter cell line was used to compare macrophage responses to two breast cancer cell types, using different types of models. For this study, we chose to use 4T1^50^ and EMT6^51,52^ cell lines, both of which represent murine triple-negative breast cancer (TNBC) and are widely used in cancer research. Despite sharing a TNBC background, EMT6 cells are considered to be less aggressive than 4T1, which are highly invasive.^53^ This suggests that these two TNBC cell lines may affect macrophages differently. First, we compared two experimental formats with each cell line: conditioned media (CM) and two-dimensional (2D) mono-layer co-culture. While CM has been used to study macrophage responses, they are not only influenced by cytokines secreted by cancer cells (found in the CM), but also by cell-to-cell interactions and the hypoxic core that forms within the tumor.^54,55^ The use of the macrophage reporter cell line allows us to directly compare simple with more complex experimental models.

The effects of 4T1 and EMT6 cell-derived conditioned media were compared to cancer cell:macrophage co-cultures generated in a 1:1 ratio. Since studies suggest that cancer cells can switch macrophages to the wound-repair (M2) subtype,^56^ we expected to see diminished fluorescence of the RAW:*iNos*-eGFP reporter following exposure to the cancer cells and CM. Our results indeed showed that the macrophage cells had statistically significant lower levels of fluorescence compared to non-treated macrophages, similarly to the M2-like phenotype, after 48 h of exposure to either conditioned media or the 4T1 or EMT6 cells (**Figure 2**). While both EMT6 models, CM and co-culture, resulted in similar effects, there was a significant difference between them (p < 0.01), with co-culture of the cells having a greater reduction in fluorescence. The 4T1 models’ outcomes were substantially different from one another. While the 4T1-CM had a statistically significant change versus non-treated cells (p < 0.001), it resulted in the least change from the non-treated control overall. On the other hand, the 4T1 co-culture yielded the lowest levels of florescence in the assay. Taken together, the 2D co-cultures produced greater effects than their CM counterparts, however, the trends observed were unexpectedly inconsistent – while co-culture with 4T1 cells yielded the lowest fluorescence, as expected, the CM from EMT6 cells resulted in greater effects than that from 4T1s. These results show the sensitivity of the reporter and highlight the relevance of cell-to-cell interactions.

**Figure 2.**
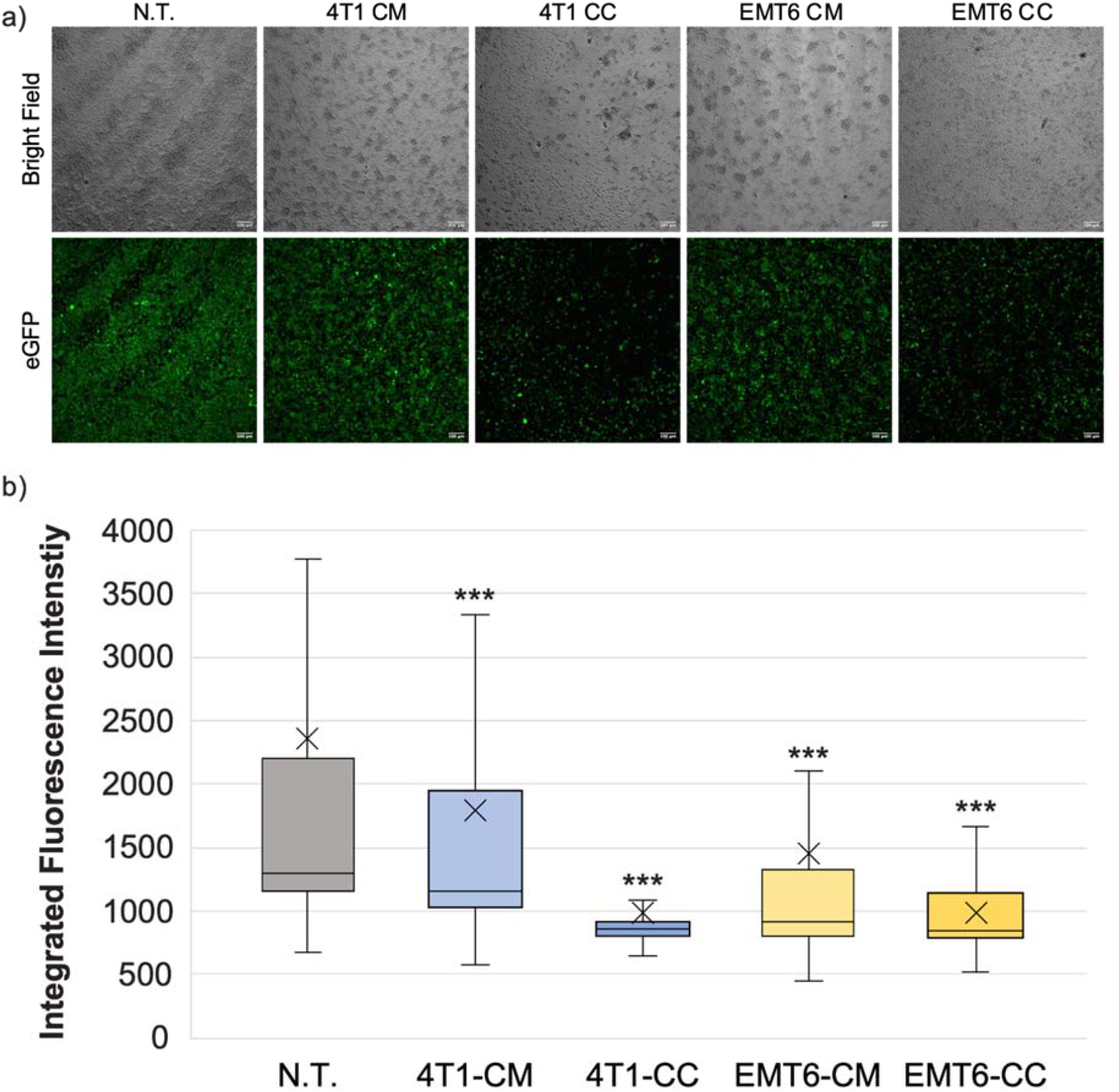
Macrophage exposure to conditioned media (CM) from or co-culture (CC) with 4T1 or EMT6 cells. (a) Confocal microscopy images of RAW:iNos-eGFP cells following exposure to conditioned media from 4T1 or EMT6 cells or subjected co-culturing with each. eGFP=enhanced green fluorescent protein. Scale bars on confocal images represent 100 µm. (b) Per-cell fluorescence intensity quantification of images shown in (a). The “x” in the box represents the mean; the bottom and top lines of the box represent the median of the bottom half (1st quartile) and median of the top half (3rd quartile), respectively; the line in the middle of the box represents the median; the whiskers extend from the ends of the box to the minimum value and maximum value. Student’s T-test was used for statistical analysis versus N.T. (n = 1,422) (p<0.001 = ***). Student T-test comparing 4T1-CM (n = 968) versus 4T1-CC (n = 370) and EMT6-CM (n = 581) versus EMT6-CC (n = 336) shows statistical significance (p<0.001=*** and p<0.01=**, respectively). N.T. = not treated.

While 2D co-culture monolayers are simple to use and allow the study of intracellular signaling cascades and cell behavior, this approach omits the highly complex structural organization found in the three-dimensional TME.^57^ Also, solid tumors develop an oxygen-less region within the tumor (hypoxic core), which leads to tumor progression and metastasis and promotes macrophage polarization towards the M2 phenotype.^58,59,60^ For these reasons, we assessed the impacts of 4T1 and EMT6 spheroids as *in vitro* three-dimensional (3D) models, on the macrophage reporter cell line. To better represent the TME, only spheroids that were 400 µm or more in diameter were used, since these are known to develop a hypoxic core.^61^

Macrophages exposed to 4T1 or EMT6 spheroids showed reduced levels of *iNos*-controlled eGFP expression, suggesting that they are adopting a tumor-promoting, M2 phenotype (**Figures 3** and **S4**). Interestingly, while both 4T1 and EMT6 spheroids seemed to reduce the eGFP signal, the change was greater when macrophages were exposed to 4T1 spheroids, which is consistent with the contrasting aggressiveness of the model cell lines. This also highlights the advantages of using a reporter-based approach to monitor the interactions between macrophages and cancer cells, while keeping track of the macrophage polarization state. The use of our reporter cell line not only facilitates the study of macrophage phenotypes in real time, but also allows the use of more complex and relevant cancer models that could not be used before due to experiment- or technique-associated limitations.

**Figure 3.**
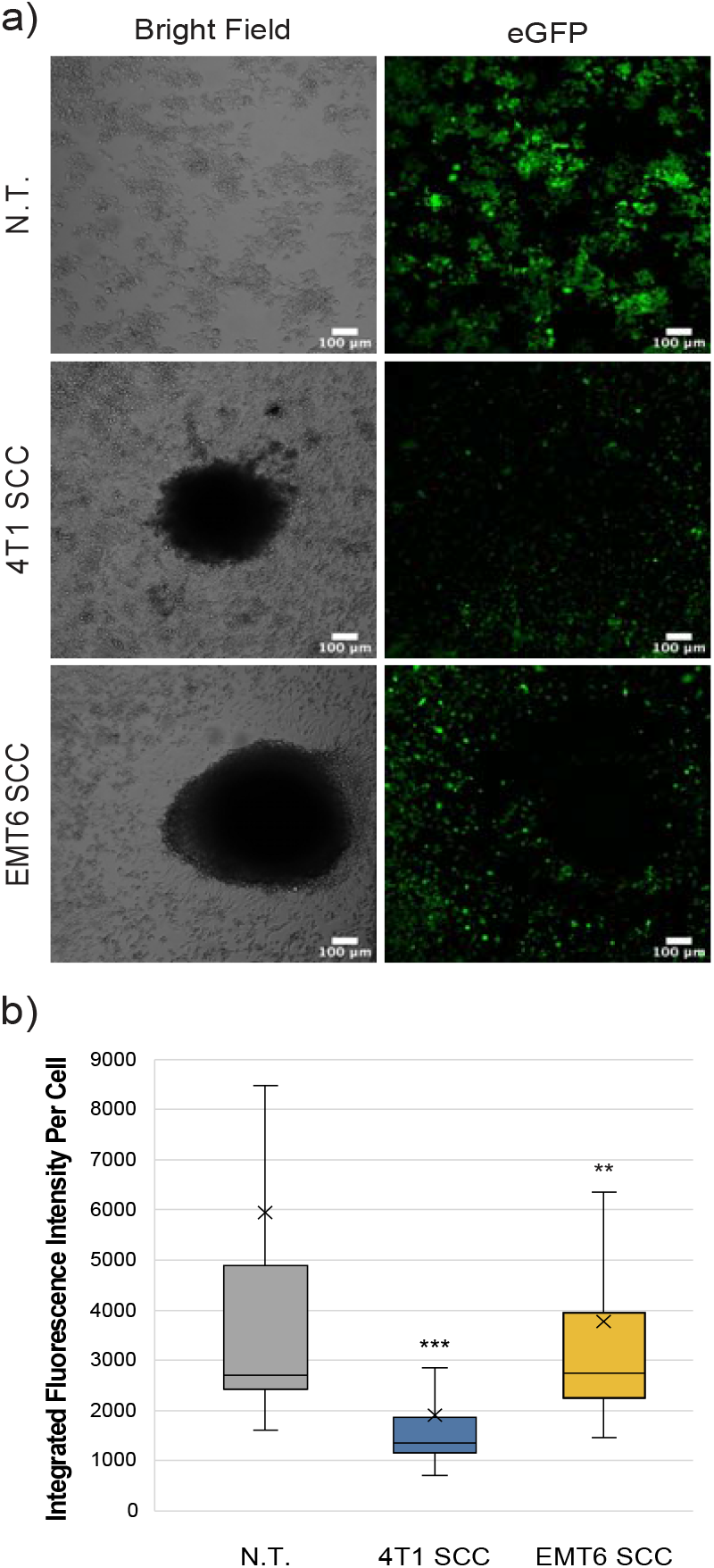
Macrophage reporter responses in three-dimensional tumor models. (a) Representative confocal images of RAW:*iNos*-eGFP macrophages (top row) after 48 h exposure to 4T1 (middle row) or EMT6 (bottom row) spheroids. eGFP = enhanced green fluorescent protein. Scale bars represent 100 µm. Additional spheroids are shown in **Figure S4**. (b) Per-cell fluorescence quantification of confocal images from (a). The “x” in the box represents the mean; the bottom and top lines of the box represent the median of the bottom half (1st quartile) and median of the top half (3rd quartile), respectively; the line in the middle of the box represents the median; the whiskers extend from the ends of the box to the minimum value and maximum value. Student T-test was used for statistical analysis of 4T1 SCC (n = 406) and EMT6 SCC (n = 337) versus N.T. (n = 2,111), (p<0.01 = **, p<0.001 = ***). N.T.= non-treated cells, SCC=spheroid co-culture.

### Re-Programming of RAW:*iNos*-eGFP Cells in the Presence of Spheroids via Small Molecule Treatment

To further assess the interactions of macrophages with breast cancer and evaluate their reprogramming capabilities in tumor microenvironments, we used pyrimido(5,4-b)indole (PBI1), a Tlr-4 agonist known to activate macrophages to M1-like phenotypes and enhance their anti-cancer activity.^6^ First, we tested the effects of different PBI1 concentrations (5, 10, and 20 µg/mL) on the viability of RAW:iNos-eGFP cells using Alamar Blue reagent and compared it to non-treated and 0.4% DMSO-treated control cells; no substantial changes were observed (**Figure S5a**). RT-PCR quantifying *iNos* mRNA transcript levels was also performed to confirm the activation of macrophages by PBI1 (**Figure S5b**), as was an assessment of fluorescence changes in the RAW:*iNos*-eGFP cells (**Figure 4**). In both cases, PBI1 treatment resulted in expected increases relative to the control groups, confirming that the small molecule’s promotion of macrophage activation persists in the reporter cells.

**Figure 4:**
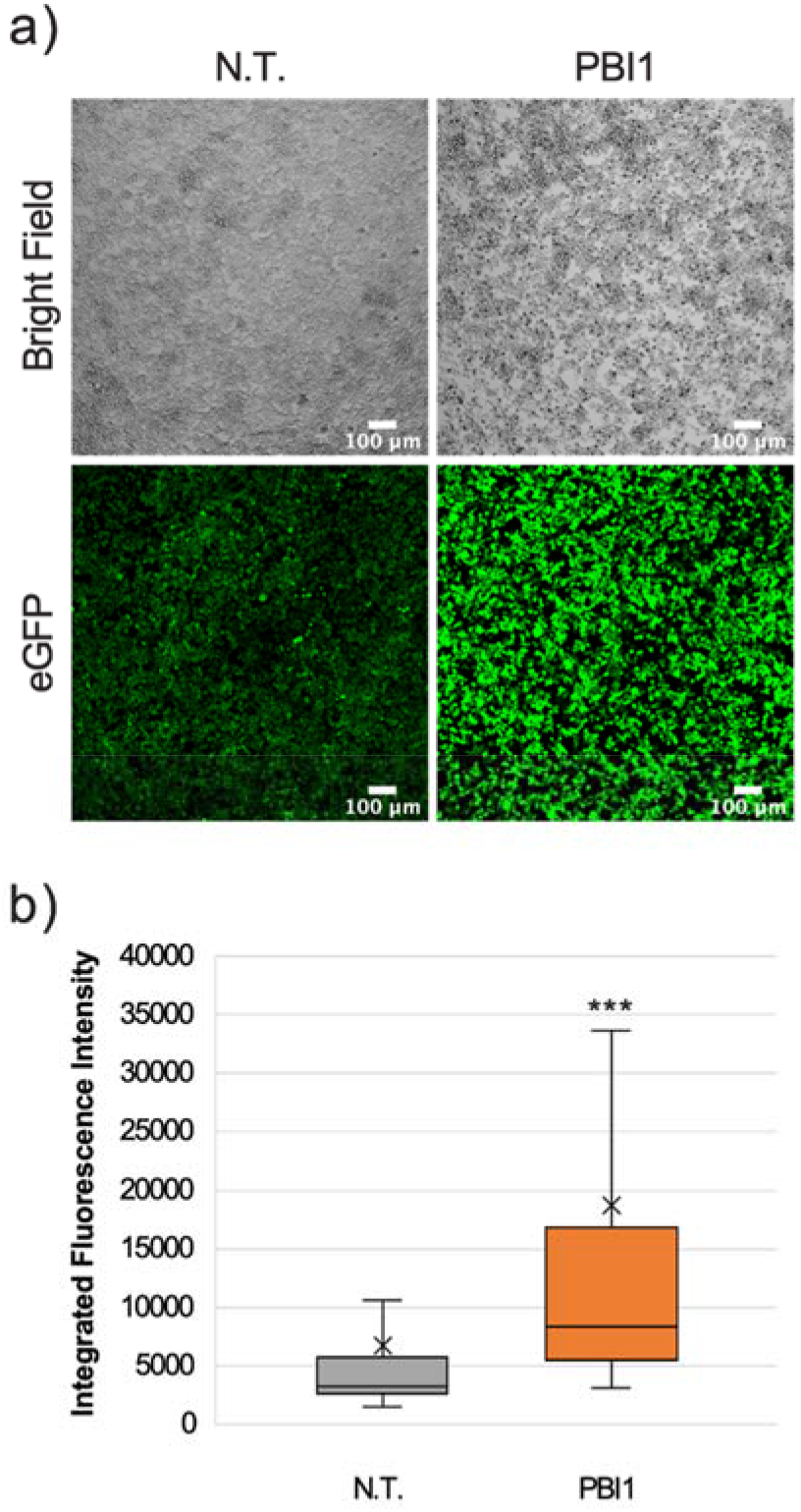
Effects of PBI1 on RAW:iNos-eGFP cells. (a) Confocal images showing changes in fluorescence following PBI1 treatment. Scale bars represent 100 µm. (b) Quantification of fluorescence intensity from images in (a). The “x” in the box represents the mean; the bottom and top lines of the box represent the median of the bottom half (1st quartile) and median of the top half (3rd quartile), respectively; the line in the middle of the box represents the median; the whiskers extend from the ends of the box to the minimum value and maximum value. Error bars represent standard error. Student T-test was performed versus N.T. (p<0.001 = ***). N.T.=non-treated cells, eGFP=enhanced green fluorescent protein.

We then used PBI to affect RAW:*iNos*-eGFP macrophages co-cultured with either 4T1 or EMT6 spheroids; controls lacked treatment with the small molecule. As in our earlier experiment (**Figure 3**), reporter macrophages exposed to either 4T1 or EMT6 spheroids in the absence of PBI1 displayed decreased fluorescence (**Figure 5**). Excitingly, a significant increase in *iNos*-eGFP signal was observed following treatment with PBI1, suggesting a shift of the macrophages toward a pro-inflammatory, anti-tumor, M1 phenotype. While the change observed was not statistically significant, it is important to note that the mean values for 4T1 SCC (+) and EMT6 SCC (+) groups increased relative to the N.T. (-) group. We also investigated the effects of PBI1 treatment on 4T1 and EMT6 spheroids in the absence of macrophages to verify that the molecule itself did not affect them (**Figure S6**). These results reinforce the idea that macrophages can be reprogrammed at the tumor site, and the reporter can be used to track these changes, even in more complex models of the TME.

**Figure 5:**
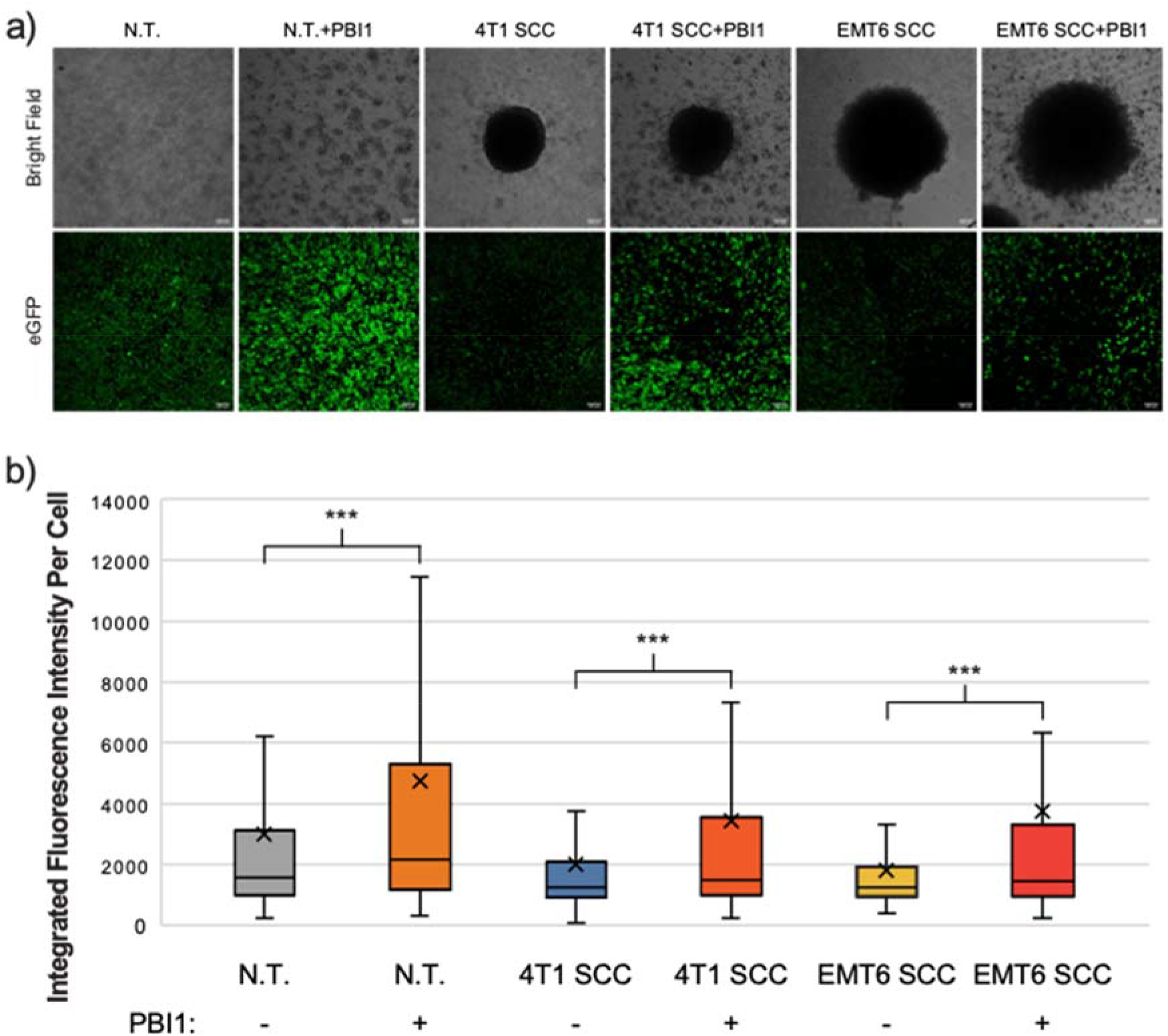
Re-polarization of RAW:iNos-eGFP macrophages using a small molecule in a spheroid co-culture model. (a) Confocal microscopy images of RAW:iNos-eGFP cells co-cultured with 4T1 or EMT6 spheroids, with (+) or without (-) PBI1 treatment. Scale bars on confocal images represent 100 µm. Images of additional spheroids are shown in **Figure S6**. (b) Student T-test was used for statistical analysis versus PBI1-treated groups (p<0.001 = ***). N.T. (-): n = 2991; N.T. (+): n = 1536; 4T1 SCC (-): n = 703; 4T1 SCC (+): n = 539; EMT6 SCC (-): n = 572; EMT6 SCC (+): n = 276. The “x” in the box represents the mean; the bottom and top lines of the box represent the median of the bottom half (1st quartile) and median of the top half (3rd quartile), respectively; the line in the middle of the box represents the median; the whiskers extend from the ends of the box to the minimum value and maximum value. N.T. = non-treated, SCC = spheroid co-culture.

## 4. Discussion

In summary, our study demonstrates the advantages of using a macrophage phenotype-reporter cell line, RAW:*iNos*-eGFP to study macrophage interactions with cancer, especially in more complex environments. Following the confirmation of a consistently expressed, phenotype-specific marker, *iNos* (**Figure S2**), we designed a cell line based on the commonly used macrophage model, RAW264.7 cells, to express eGFP under the regulation of the *iNos* promoter. After validating normal functioning of the cells via RT-PCR and reporter fidelity via confocal microscopy (**Figure 1**), we explored the responses of macrophages to different breast cancer models derived from 4T1 and EMT6 TNBC cell lines.

Across the three models examined, conditioned media (CM), 2D co-culture (CC), and 3D/spheroid co-cultures (SCC), both TNBC cell lines resulted in anti-inflammatory M2-like fluorescence profiles (**Figures 2** and **3**). However, while we expected the 4T1-derived models to demonstrate greater changes than those from the less aggressive EMT6 cells, this was not reflected in the conditioned media experiments. For CM, EMT6 gave a greater response than 4T1; the CC and SCC models showed the opposite -- the 4T1 cells resulted in greater changes to *iNos*-driven eGFP expression than EMT6. These results confirm that there are differences in macrophage responses among models used, even with the same cell line(s), and should be considered in future studies.

The re-education of macrophages in disease states or infections, either from pro- to anti-inflammatory or vice-versa, is a therapeutic strategy of broad interest.^62^ In cancer, the polarization of macrophages to the immune-stimulating (M1) phenotype can result in enhanced anti-tumor activity, and is a goal in developing new cancer immunotherapies.^23,63,64^ Therefore, having assessed the RAW:*iNos*-eGFP reporters’ functioning in TME models, we decided to test our platform in tracking macrophage re-education in the most disease representative model used here, spheroid co-culture. We used a previously described small molecule Tlr-4 agonist, PBI1.^6^ This compound had been shown to induce M1-like polarization and enhance macrophage anti-cancer activity *in vitro*, so we put it to test in the presence of more relevant 3D tumor models. We generated spheroids using 4T1 and EMT6 cells and using only those that were 400 µm or more in diameter, compared non-treated 4T1 or EMT6 spheroid co-cultures with PBI1-treated groups. As expected, the controls (macrophages exposed to spheroids without PBI1), exhibited decreased fluorescence representative of an anti-inflammatory, M2-like phenotype. However, when challenged with the Tlr-4 agonist small molecule, PBI1, macrophages showed increases in fluorescence relative to the non-treated group, indicative of their activation toward a pro-inflammatory state. This illustrates the feasibility of using reporters to evaluate therapeutics even in the presence of more relevant and realistic cancer TME models.

In conclusion, we show that the use of the macrophage reporter cell line enables the assessments across different models without needing to apply other probes (e.g., antibodies) for detection, which unless used with fixed cells, could alter cellular characteristics, or segregate or lyse cells. This approach also allows for continuous monitoring of macrophage responses, as conditions change or additional stimuli are presented. Taken together, the use of macrophage reporter cell lines, including the one developed here or others with suitable phenotype-specific markers, can facilitate studies to assess macrophage behavior in response to more complex and accurate models, and high-throughput assessment of drugs for affecting macrophages.

## Supporting information

Supporting Information for Mas-Rosario et al.

## 6. Author Contributions

JAM-R. and MEF conceived the research study. JAM-R planned the experiments, JAM-R, JDM, MIJ, and JM-M performed the experiments and conducted data analysis. JAM-R and MEF interpreted the results obtained. JAM-R. wrote the manuscript with assistance from MEF. All authors approved the final manuscript.

## 8. Acknowledgements

Thanks to Prof. Scott Garman (Biochemistry & Molecular Biology, UMass Amherst) for helpful input on this project, and Prof. D. Joseph Jerry (Veterinary and Animal Sciences, UMass Amherst) for supplying materials associated with stable transfections. Special thanks to the Vishnu Raman, Ph.D., Shoshanna Bloom, Ph.D., and Victoria Wetherby (Chemical Engineering, UMass Amherst) for sharing their knowledge of molecular cloning and Neeraj Raghuraman and Yasushi Kimura, M.D. (Mechanical Engineering, UMass Amherst) for assistance in confocal imaging and fluorescence quantification. Flow cytometry and cell sorting were performed at the University of Massachusetts Amherst Institute for Applied Life Sciences (IALS) Flow Cytometry Core Facility; we thank facility staff Dr. Amy Burnside and Arsh Patel for their assistance.

