## Supporting Information for Mas-Rosario et al. for "Macrophage-Based iNos Reporter Reveals Polarization and Reprogramming in the Context of Breast Cancer"

Supplementary Material

### 1. **Materials and Methods**

##### **Isolation and differentiation of primary macrophages from bone marrow**

Isolation and differentiation of primary macrophages was performed following a previously established procedure.^1^ Bones were collected from femurs and tibiae from C57/BL6 mice and put into 0.6 mL micro-centrifuge tubes containing a small hole at the bottom (perforated prior to use with an 18G needle). Each 0.6 mL micro-centrifuge tube (containing one femur and one tibia) was then inserted into a 1.5 mL micro-centrifuge tube and centrifuged for 30 seconds at 10,000 rpm. The 0.6 mL tube with marrow-less bones was discarded and the pelleted bone marrow (now in the 1.5 mL tube) was re-suspended in 500 µL media containing 70% complete DMEM and 30% L929-conditioned media (referred to as differentiation media). L929 cells were obtained from Professor Barbara Osborne (Veterinary and Animal Sciences, UMass Amherst), and cultured and used to generate conditioned media using the same procedures as described in the main text. The contents of each 1.5 mL tube were then transferred to a T175 tissue culture flask, mixed with 12 mL of differentiation media, and grown for 3 days. After 3 days, the media was replaced with 12 mL of fresh differentiation media for another 4 days. Following differentiation, media was replaced with complete DMEM and the cells were kept in T175 culture flasks at a density of 1-2 x 10^6^cell/mL at 37 °C under a humidified atmosphere containing 5% CO_2_ until needed for experiments (for a maximum of 3 weeks).

2. **Supplementary Figures**

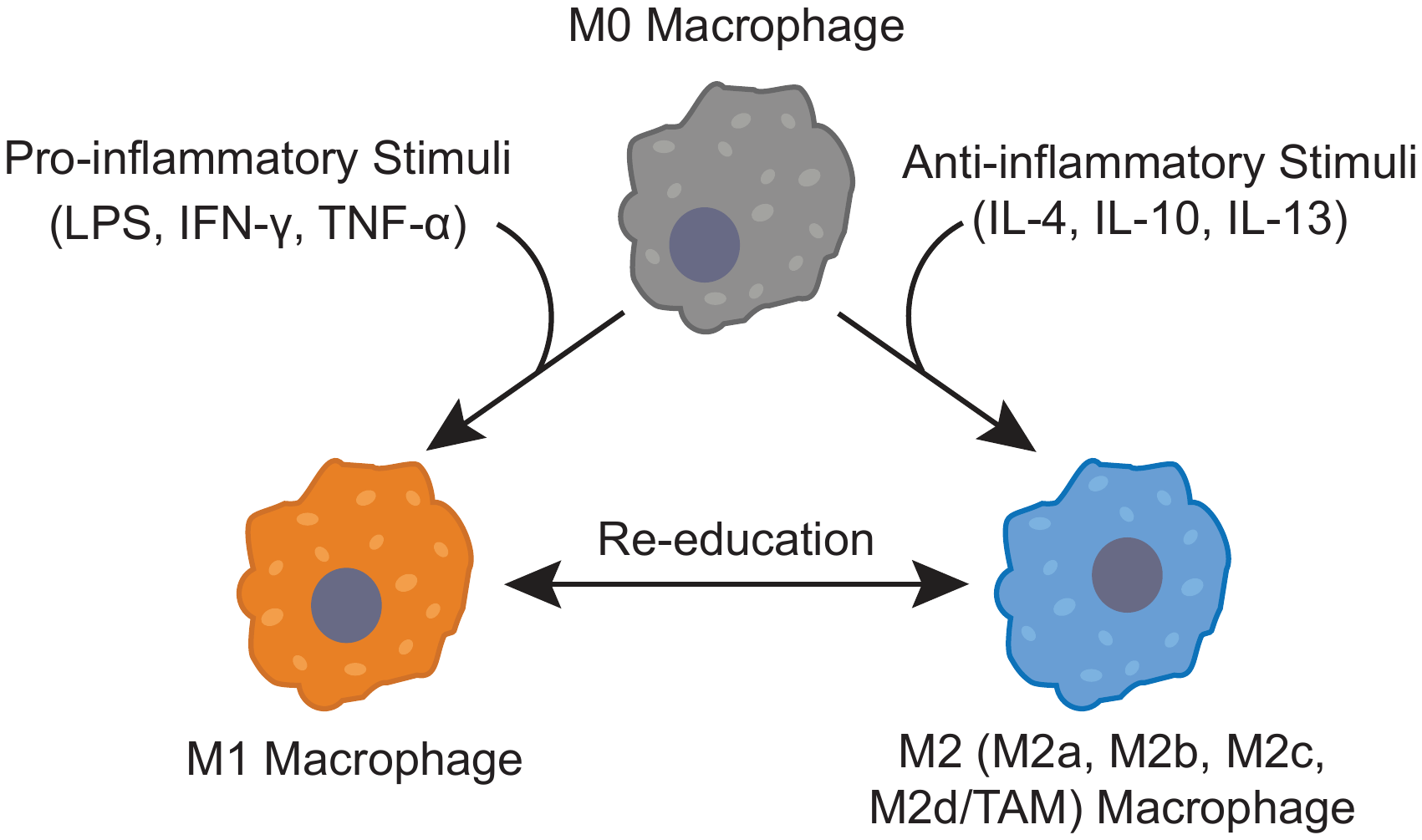

**Figure S1: Macrophage polarization.** From an undifferentiated, naïve (M0) state, macrophage cells assume a pro-inflammatory (M1) phenotype (lower left, orange) when activated with pro-inflammatory cytokines (e.g., lipopolysacharide (LPS), interferon gamma (IFN-γ), and/or tumor necrosis factor alpha (TNF-α)). When stimulated with anti-inflammatory cytokines such as interleukin 4 (IL-4), interleukin 10, (IL-10), or interleukin 13 (IL-13), macrophages are converted into an anti-inflammatory (M2) subtype (lower right, blue). M2 macrophages have multiple sub-classifications (M2a, M2b, M2c, and M2d/TAMs), which result from different stimuli and have varying characteristics. Once polarized, macrophages can be “re-educated” to a different phenotype, e.g., M2 to M1. TAM = tumor associated macrophage.

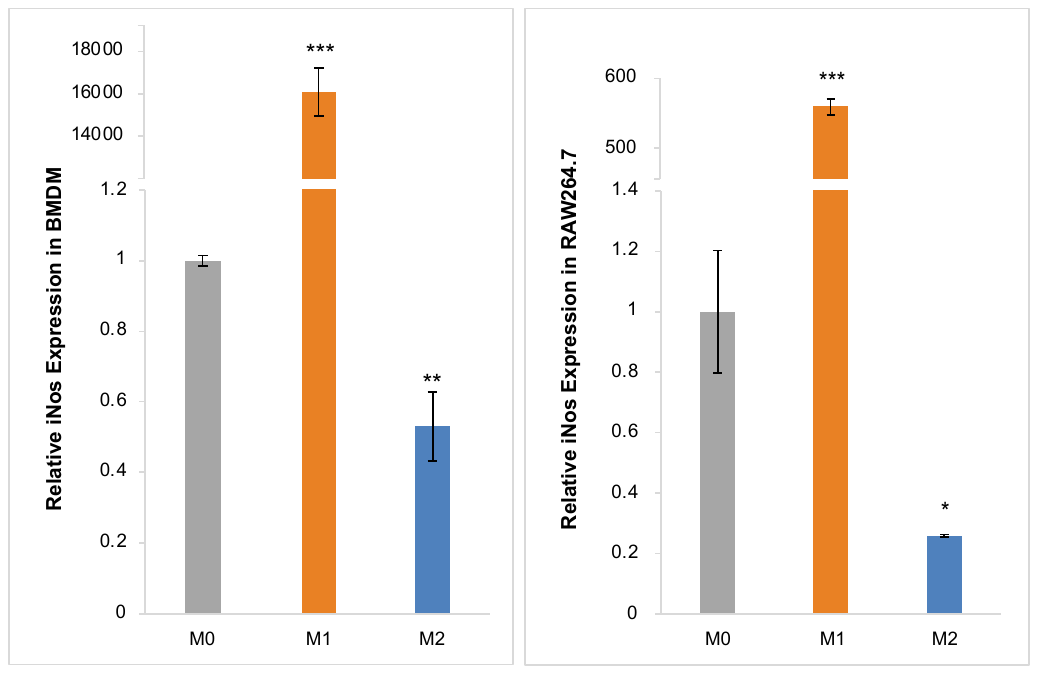

**Figure S2: RT-PCR analysis comparing *iNos* levels in immortalized versus primary macrophages.** Shown are relative levels of *iNos* expression in bone marrow-derived macrophages (BMDMs; left) versus RAW264.7 (right) cells under polarizing conditions. M0 (gray)=non-treated, M1 (orange)=LPS/IFN-γ, and M2=IL-4. This experiment was performed with three biological replicates. Student T-test was used for statistical analysis (p<0.05 = *, p<0.01 = **, p<0.001 = ***). Error bars represent standard error.

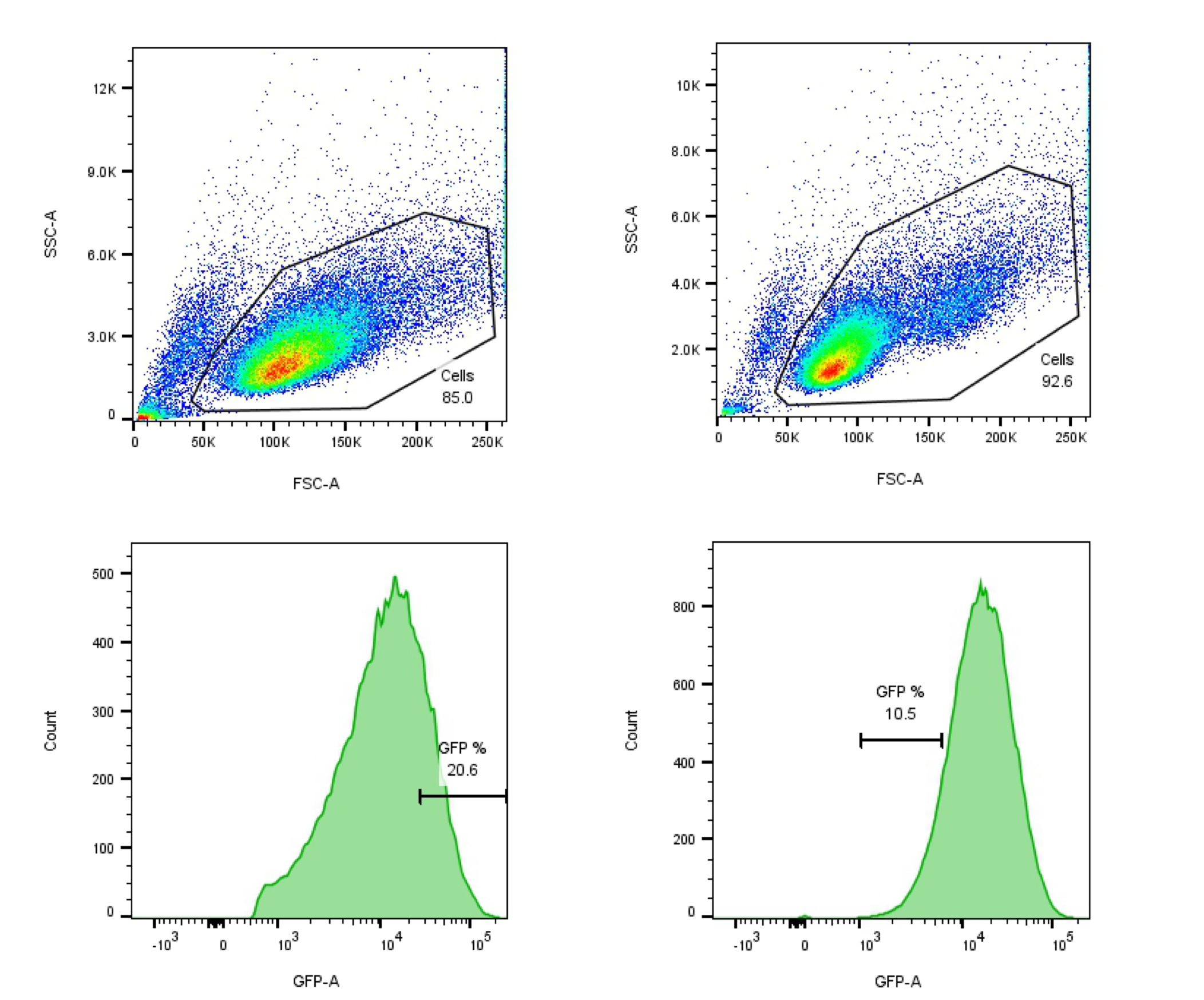

**Figure S3: Sorting of RAW:iNos-eGFP cells.** Cytometry plots from the first (top and bottom left) and second (top and bottom right) sortings of RAW:*iNos*-eGFP cells under M1- followed by M2-polarizing conditions, respectively. For the first sorting under M1 conditions (top and bottom left), selected GFP-positive cells are encased (top left image), from the top 20.6% of positive cells were kept (bottom left image). The cells were then sorted again under M2 conditions (top and bottom right), and the bottom 10.5% of positive cells were kept.

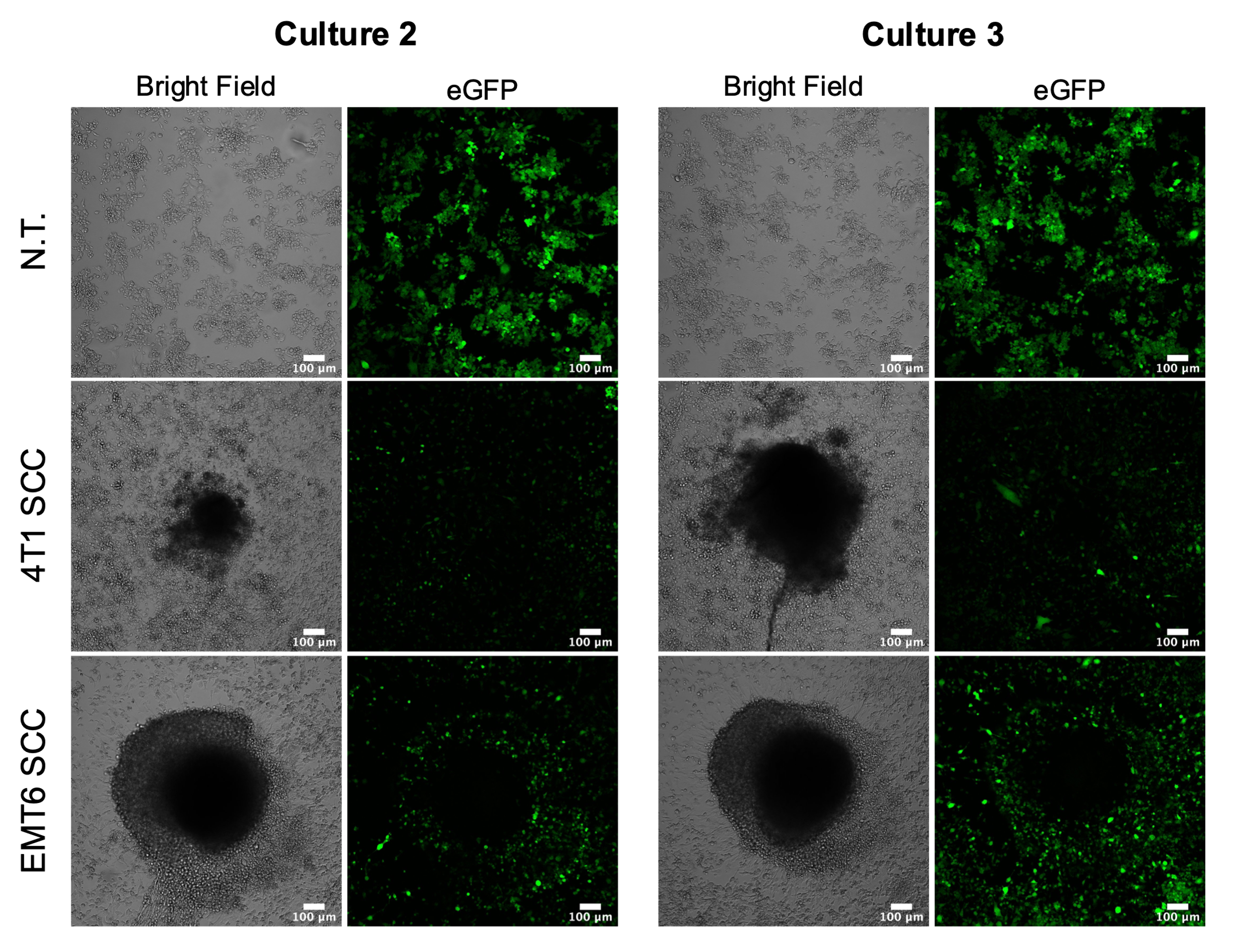

**Figure S4: Additional 3D co-cultures.** To supplement images shown in **Figure 3**, representative images from additional cultures/spheroids are shown here. RAW:*iNos*-eGFP macrophages (top row) are shown after 48 h exposure to 4T1 (middle row) or EMT6 (bottom row) spheroids. eGFP = enhanced green fluorescent protein. Scale bars represent 100 µm.

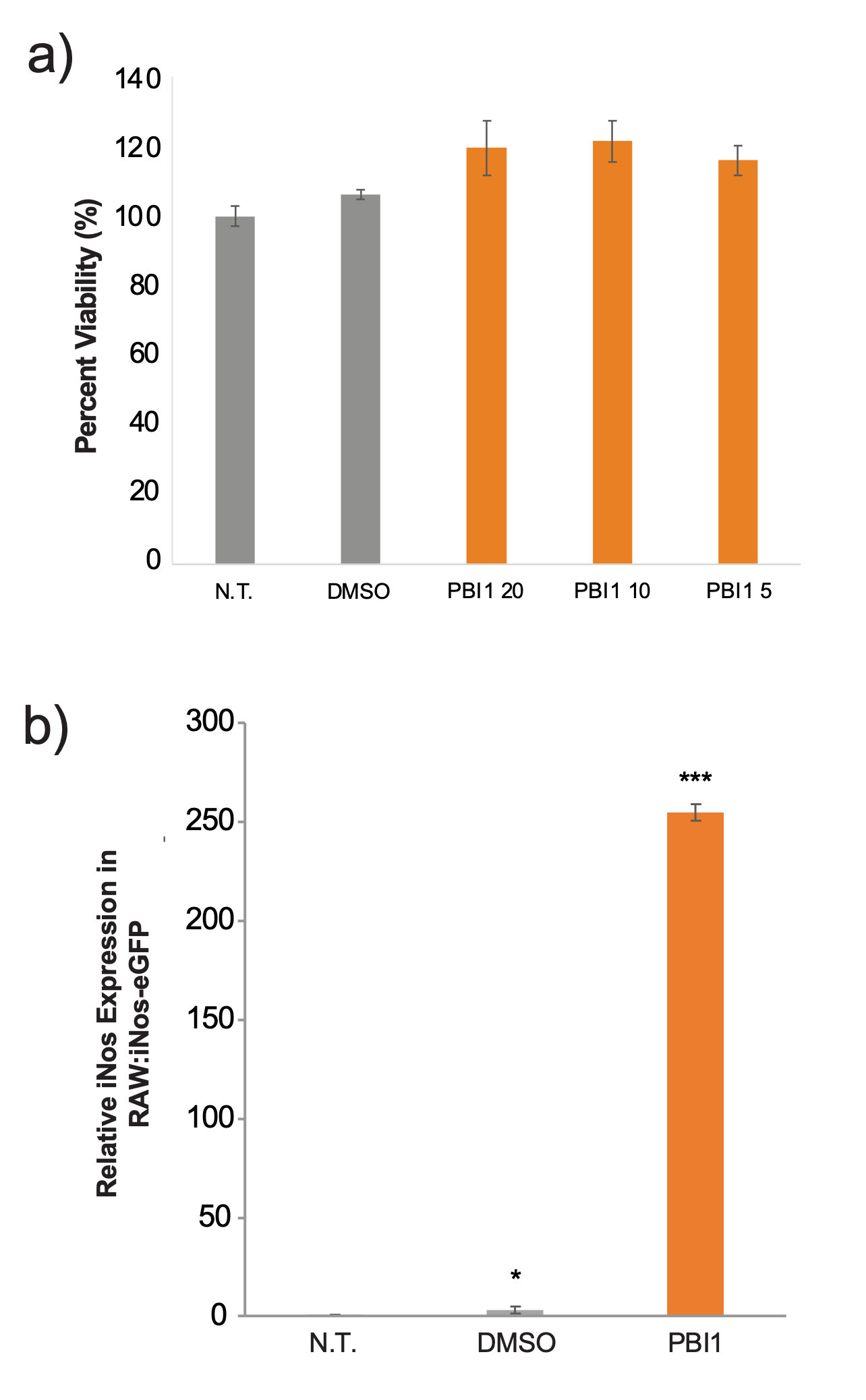

**Figure S5: Evaluation of PBI1 effects on RAW:*iNos*-eGFP cells.** (a) Viability assay of PBI1-treated cells using three different concentrations (5, 10, and 20 µg/mL). (b) RT-PCR results showing relative *iNos* mRNA transcript levels in RAW:*iNos*-eGFP cells following PBI1 treatment at a final concentration of 20 µg/mL. Student T-test comparing DMSO- and PBI1-treated groups to N.T. (p<0.05 = *, p<0.001 = ***). N.T. = non-treated cells, DMSO = cells treated with 0.4% dimethyl sulfoxide vehicle. Error bars represent standard error.

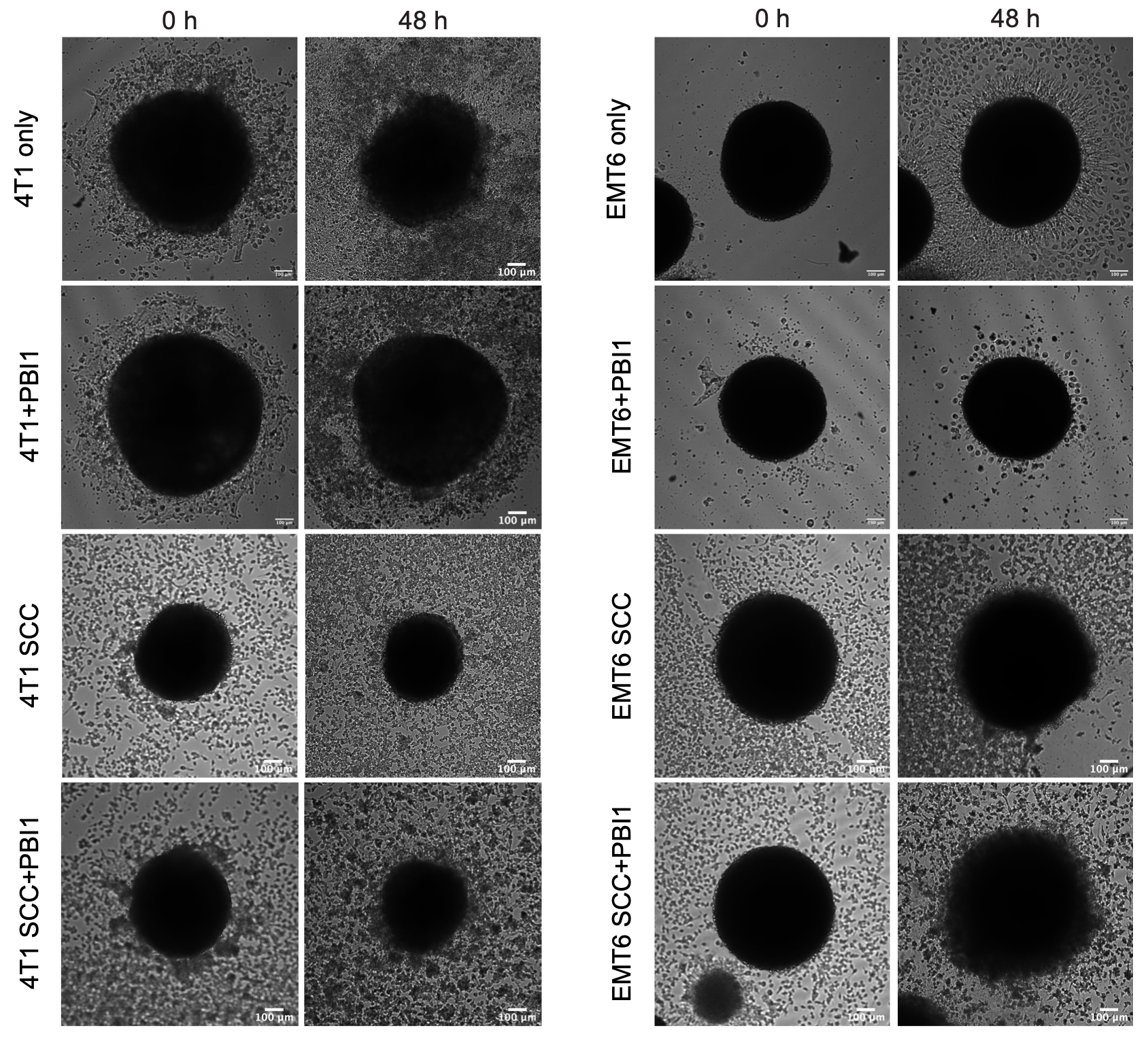

**Figure S6: Effects of PBI1 on 4T1 and EMT6 spheroids after 48 h.** Confocal microscopy images showing changes in spheroid size after 48 h with or without treatment with 20 µg/mL of PBI, and spheroid co-culture with RAW:iNos-eGFP cells (SCC). Scale bars on images represent 100 µm.

3. **References**

1. Weischenfeldt J, Porse B. Bone Marrow-Derived Macrophages (BMM): Isolation and Applications. *Cold Spring Harb Protoc* (2008) pdb.prot5080. doi: 10.1101/pdb.prot5080.
